## Supplementary material for "The molecular basis of Human FN3K mediated phosphorylation of glycated substrate": Supplmental Information

**Figure S1-**

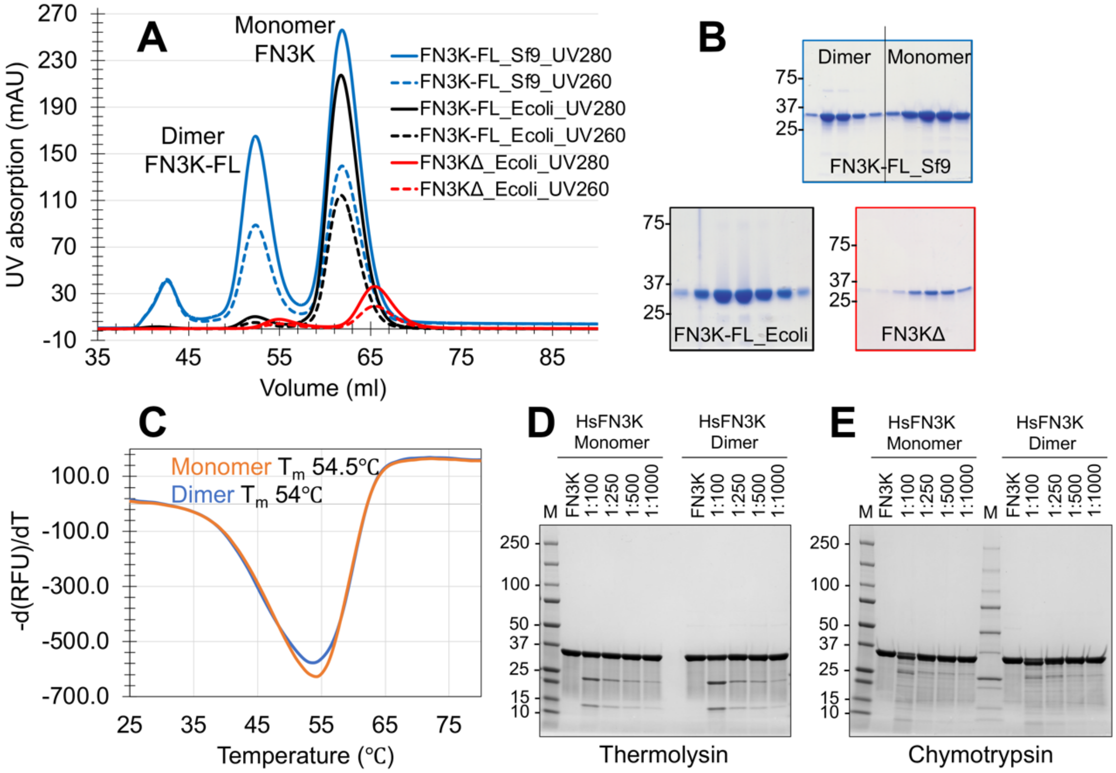


**Figure S1**- Purification and biophysical characterization of HsFN3K. (A) SEC chromatogram of HsFN3K purified from different expression sources. FN3K expressed in Sf9 cells showed a significant dimeric peak in SEC. (B) SDS-PAGE showing different purified HsFN3K proteins. (C) Thermal melting spectra for the dimeric and monomeric HsFN3K (expressed in Sf9) species. The melting temperature (Tm) for both species are very similar, suggesting a similar structural fold for both. Limited proteolysis of HsFN3K (Sf9) monomer and dimer using (D) Thermolysin and (E) Chymotrypsin proteases. The w/w ratio of protein to protease is noted at the top of each lane. “M” is a protein ladder.

**Figure S2-**

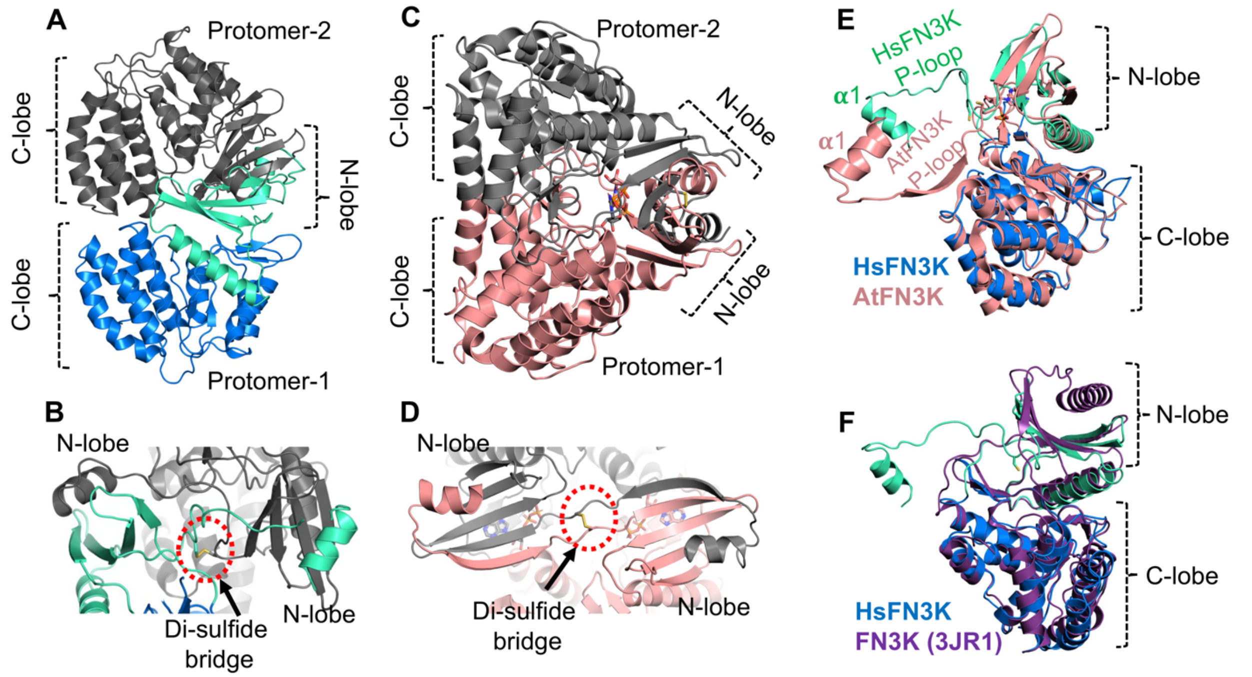


**Figure S2-** A comparison of HsFN3K and AtFN3K crystal structures. (A) Cartoon representation of the HsFN3K dimer in the asymmetric unit in the apo-HsFN3KΔ crystal structure. (B) The P-loop Cys24 (shown in sticks) disulfide bridge in the HsFN3KΔ dimer. (C) Cartoon representation of AtFN3K dimer as observed in the asymmetric unit. (D) The P-loop Cys (shown in sticks) disulfide bridge in the AtFN3K structure. (E) Superposition of the HsFN3K and AtFN3K (PDBid 6OID) structures showing the difference in P-loop orientations. (F) Structure superposition of HsFN3K and *E. coli* FN3K (PDB 3JR1) illustrating the difference in P-loop orientation in the two structures.

**Figure S3-**

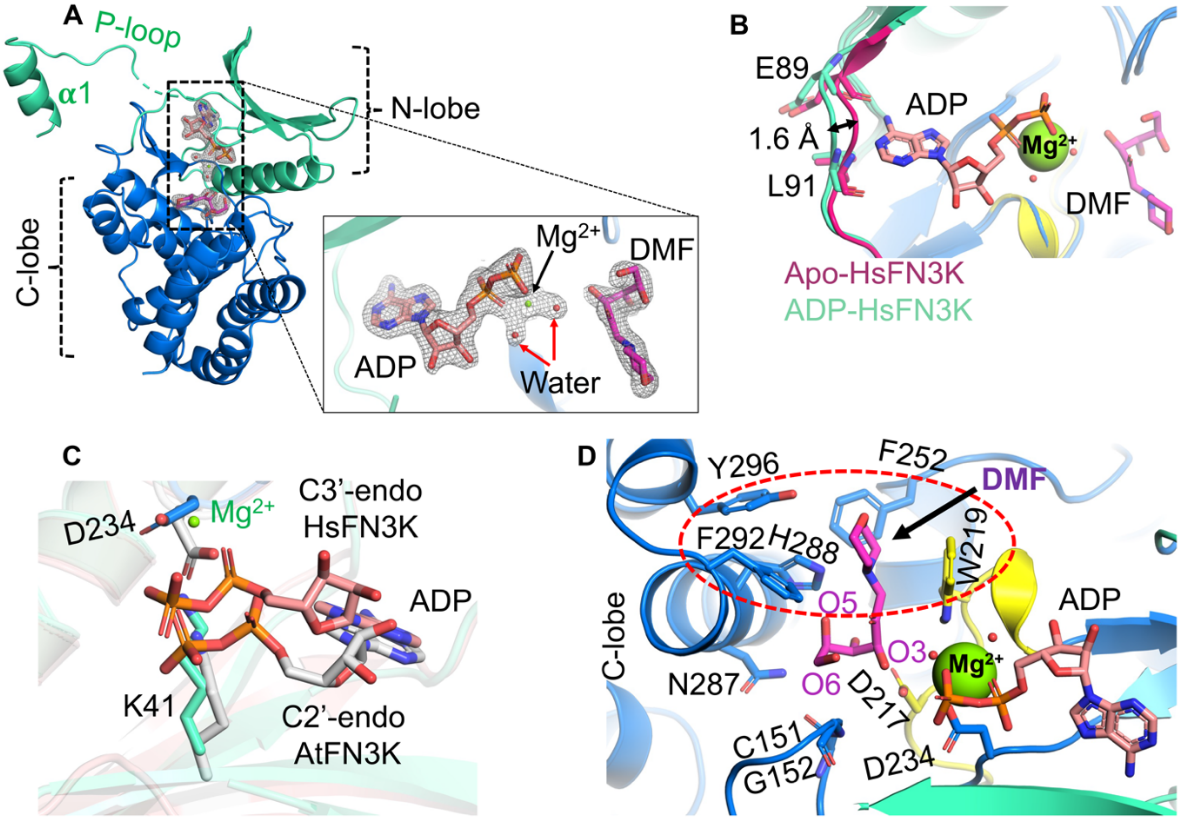


**Figure S3**- (A) Cartoon representation of the crystal structure of HsFN3K bound to ADP-DMF-$Mg^{2+}$ (protomer-1). The 2Fo-Fc electron density observed for ADP, DMF, and $Mg^{2+}$ is shown. Two water molecules coordinating the $Mg^{2+}$ ion is also visible in the electron density map. The dotted section in the P-loop represents the unmodelled region due to missing electron density (B) A close-up view of the superposed nucleotide-sensing loop in apo-HsFN3K (red) and ADP-HsFN3K (light teal) showing the subtle adjustment upon nucleotide binding. The residues involved in direct interactions with the adenine base (E89 and L91) are shown as sticks, and the loop movement is also marked. (C) ADP nucleotide binding in the HsFN3K and AtFN3K structures. The sugar geometry in HsFN3K is canonical C3'-endo, while in AtFN3K is likely in an C2'-endo conformation, resulting in a clear difference in ADP placement in the catalytic site. The C3'-endo geometry in HsFN3K favors the interaction of the nucleotide phosphates with Lys41. (D) The DMF substrate binding site in HsFN3K (shown with red dotted circle) is lined with several aromatic residues (shown as sticks). The sugar interacts directly with D217 via its phosphate-receiving O3' atom.

**Figure S4-**

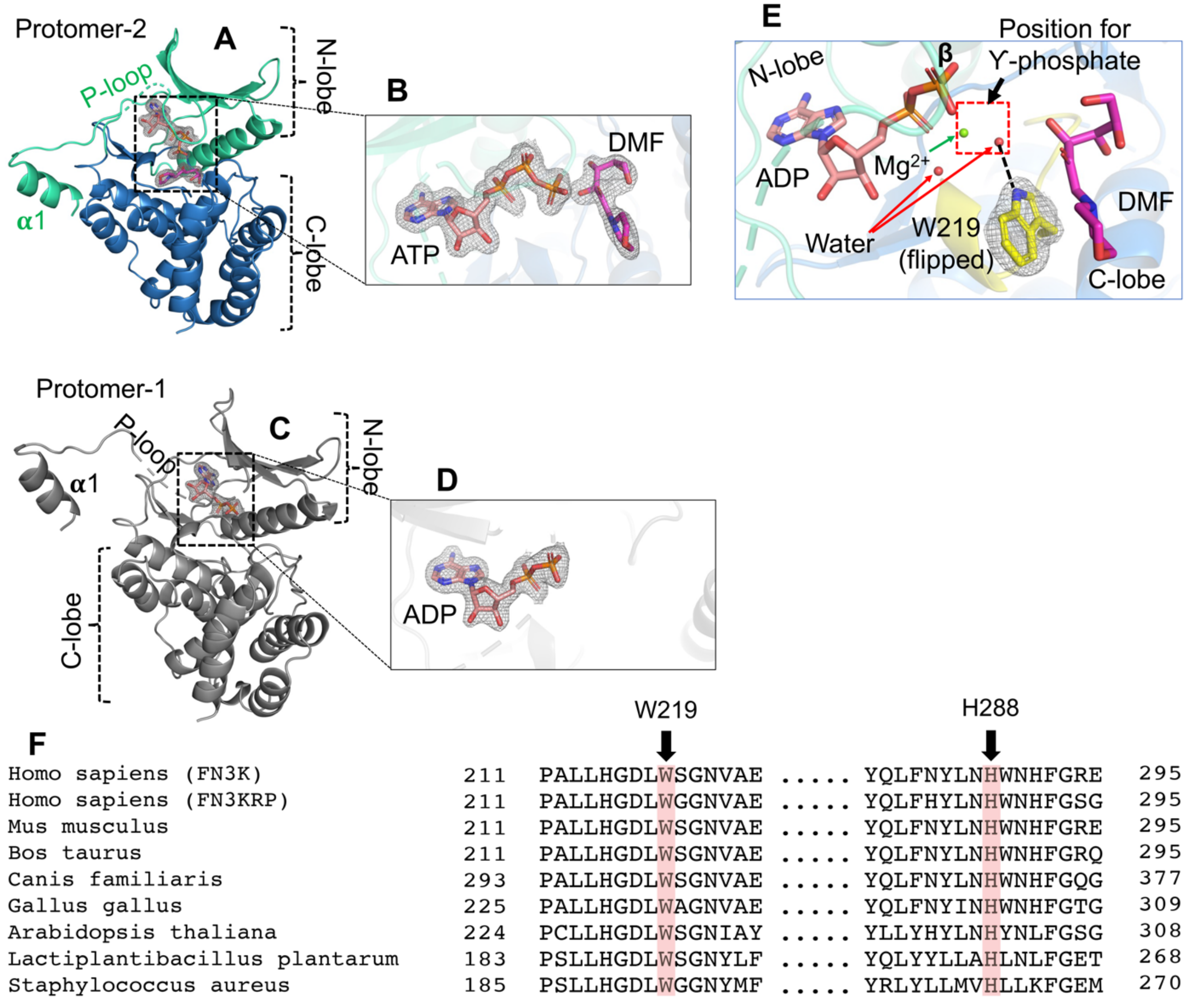


**Figure S4-** Structural features of HsFN3K bound with ATP and DMF (A) Cartoon representation of protomer-2 in the HsFN3K-ATP-DMF structure, with ATP and DMF shown as sticks. (B) 2Fo-Fc electron density for the ATP and the DMF shown at a 1σ level with a carved radius of 1.6 Å. (C) Cartoon representation of protomer-1 in HsFN3K-ATP-DMF structure, showing only ADP (in sticks) bound in the catalytic site and (D) the observed 2Fo-Fc electron density for ADP shown at a 1σ level with a carved radius of 1.6 Å. No density for DMF was observed in the substrate binding site. (E) Close-up view of the catalytic site in the HsFN3K-ADP structure showing ADP and DMF in sticks. The $Mg^{2+}$ and water molecule occupies the position of the γ-phosphate (shown as a red dotted square) which allows W219 (yellow sticks with 2Fo-Fc density shown at a 1σ level) to flip and interact with the water molecule (shown as a black dotted line). (F) The sequence alignment for FN3K shows the conservation of W219 and H288 across different species.

**Figure S5-**

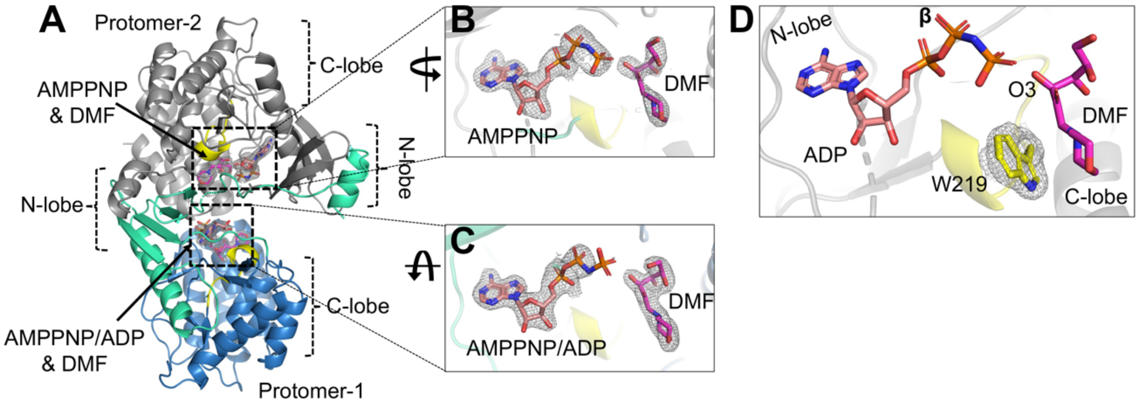


**Figure S5**- The crystal structure of HsFN3KΔ-AMPPNP-DMF shown in (A) cartoon representation. The AMPPNP and DMF observed in each protomer are shown in sticks with the superimposed 2Fo-Fc electron density shown at a 1σ level. The close-up view of the catalytic site in (B) protomer-2 and (C) protomer-1 are shown. The density for the γ-phosphate in protomer-1 is not very clear, but could still be built as AMPPNP. (D) AMPPNP binding in the protomer-2 catalytic site is unable to induce W219 flipping. The W219 is shown in predominantly the alternative conformation (as yellow sticks) with 2Fo-Fc electron density shown at 1σ level with a carved radius of 1.6 Å.

**Figure S6-**

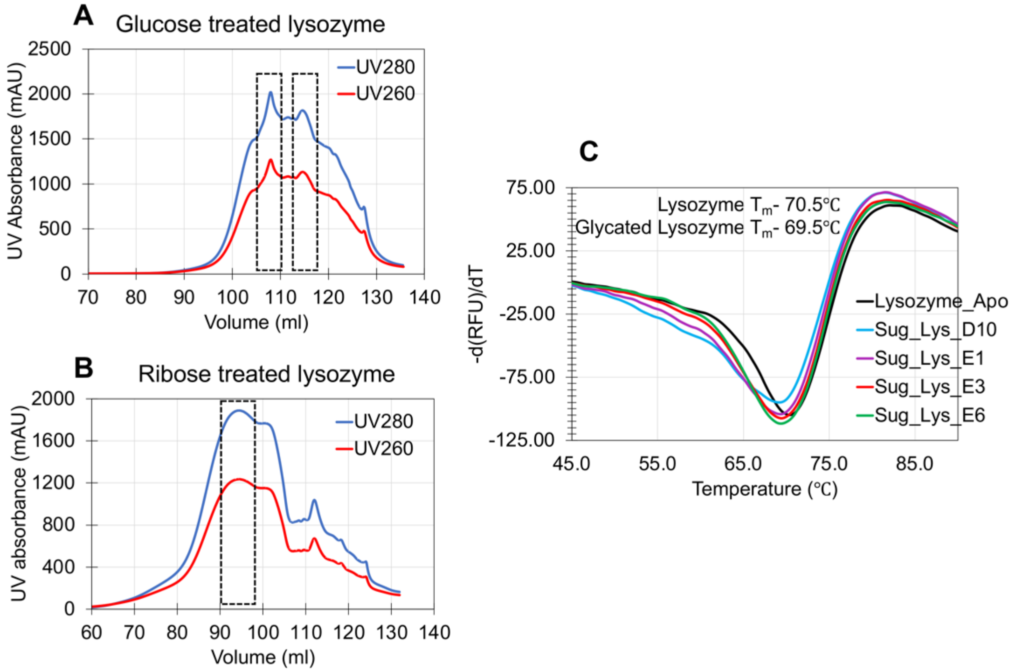


**Figure S6**- SEC chromatograms for the glycated lysozymes incubated with (A) glucose and (B) ribose sugars for 21 and 7 days, respectively. Glycated lysozyme was distributed in different peaks in both treatments and was verified by MS analysis. The dotted black box around the peaks shows the used glycated lysozyme fractions in the assays (C) The thermal melting spectra for the glycated lysozyme (with either glucose or ribose) demonstrate that the Tm following the sugar treatment did not change.

**Figure S7-**

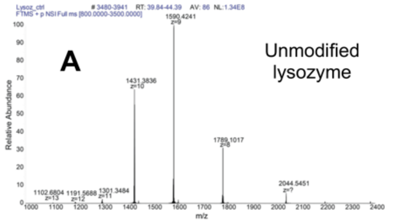


| z | Peak 1 | Glucose added | 0 |
|---|---|---|---|
| 7 | 2044.545 | | 14304.765 |
| 8 | 1789.102 | | 14304.755 |
| 9 | 1590.424 | | 14304.751 |
| 10 | 1431.384 | | 14303.763 |
| 11 | 1301.348 | | 14303.752 |
| 12 | 1191.569 | | 14286.738 |
| 13 | 1102.68 | | 14321.75 |
| | | Average (Da) | 14295.77 |

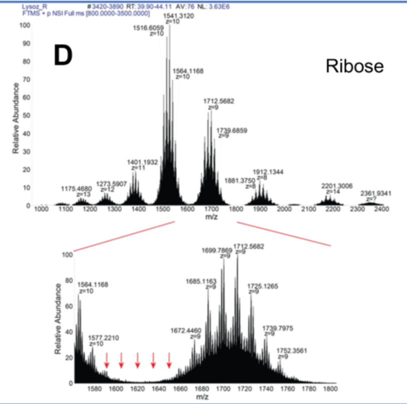


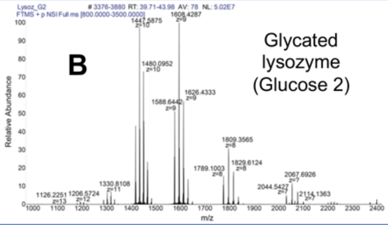


| z | Peak 1 | Peak 2 | Peak 3 | Peak 4 | Peak 5 | **Glucose added** | **0** | **1** | **2** | **3** | **4** |
|---|---|---|---|---|---|---|---|---|---|---|---|
| 7 | 2044.543 | 2067.693 | 2090.7 | 2114.136 | 2137.287 | | 14304.75 | 14466.8 | 14627.85 | 14791.9 | 14953.96 |
| 8 | 1789.1 | 1809.357 | 1829.612 | 1849.744 | 1870.001 | | 14304.74 | 14466.79 | 14628.84 | 14789.89 | 14951.95 |
| 9 | 1590.424 | 1608.429 | 1626.433 | 1644.439 | 1662.334 | | 14304.75 | 14466.79 | 14628.83 | 14790.88 | 14951.94 |
| 10 | 1431.483 | 1447.687 | 1463.891 | 1480.095 | 1496.301 | | 14304.76 | 14466.8 | 14628.83 | 14790.88 | 14952.94 |
| 11 | 1301.438 | 1316.17 | 1330.811 | 1345.543 | 1360.275 | | 14304.74 | 14466.79 | 14627.84 | 14789.89 | 14951.94 |
| 12 | 1192.985 | 1206.572 | 1220.077 | 1233.665 | 1247.086 | | 14303.73 | 14466.78 | 14628.84 | 14791.89 | 14952.94 |
| 13 | 1101.371 | 1113.836 | 1126.225 | 1138.691 | | | 14304.73 | 14466.78 | 14627.83 | 14789.88 | |
| | | | | | | Average (Da) | 14295.73 | 14457.79 | 14619.83 | 14781.87 | 14943.93 |
| | | | | | | **Δ-mass (Da)** | **0** | **162.06** | **162.04** | **162.04** | **162.06** |

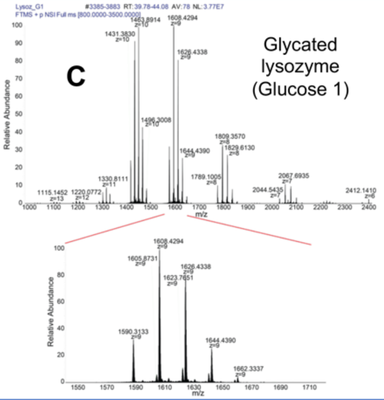


| z | Peak 1 | Peak 2 | Peak 3 | Peak 4 | Peak 5 | **Glucose added** | **0** | **1** | **2** | **3** | **4** |
|---|---|---|---|---|---|---|---|---|---|---|---|
| 7 | 2044.545 | 2067.694 | 2090.7 | 2113.993 | 2137.287 | | 14304.77 | 14466.8 | 14627.85 | 14790.9 | 14953.96 |
| 8 | 1789.101 | 1809.357 | 1829.613 | 1849.744 | 1870.251 | | 14304.75 | 14466.8 | 14628.85 | 14789.9 | 14953.95 |
| 9 | 1590.313 | 1608.429 | 1626.434 | 1644.439 | 1662.334 | | 14303.75 | 14466.8 | 14628.84 | 14790.89 | 14951.94 |
| 10 | 1431.383 | 1447.688 | 1463.892 | 1480.096 | 1496.301 | | 14303.76 | 14466.81 | 14628.84 | 14790.88 | 14952.94 |
| 11 | 1301.347 | 1316.079 | 1330.811 | 1345.543 | 1360.548 | | 14303.74 | 14465.79 | 14627.84 | 14789.89 | 14954.95 |
| 12 | 1192.984 | 1206.656 | 1220.077 | 1233.665 | 1246.919 | | 14303.73 | 14467.79 | 14628.84 | 14791.89 | 14950.94 |
| 13 | 1101.371 | 1113.837 | 1126.302 | 1138.768 | 1151.234 | | 14304.73 | 14466.78 | 14628.84 | 14790.88 | 14952.94 |
| | | | | | | Average (Da) | 14295.73 | 14457.8 | 14619.84 | 14781.88 | 14943.93 |
| | | | | | | **Δ-mass (Da)** | **0** | **162.07** | **162.04** | **162.04** | **162.05** |

**Figure S7-** The intact mass Mass-Spectrophotometry (MS) analysis for (A) unmodified lysozyme and (B-C) glucose glycated lysozymes. Thermo QualBrowser calculated mass difference (Δmass) and added glucose molecules are listed in respective tables next to the spectra (D) Intact MS analysis for ribose-modified lysozyme.
